# Vaccination with SARS-CoV-2 Spike Protein and AS03 Adjuvant Induces Rapid Anamnestic Antibodies in the Lung and Protects Against Virus Challenge in Nonhuman Primates

**DOI:** 10.1101/2021.03.02.433390

**Authors:** Joseph R. Francica, Barbara J. Flynn, Kathryn E. Foulds, Amy T. Noe, Anne P. Werner, Ian N. Moore, Matthew Gagne, Timothy S. Johnston, Courtney Tucker, Rachel L. Davis, Britta Flach, Sarah O’Connell, Shayne F. Andrew, Evan Lamb, Dillon R. Flebbe, Saule T. Nurmukhambetova, Mitzi M. Donaldson, John-Paul M. Todd, Alex Lee Zhu, Caroline Atyeo, Stephanie Fischinger, Matthew J Gorman, Sally Shin, Venkata Viswanadh Edara, Katharine Floyd, Lilin Lai, Alida Tylor, Elizabeth McCarthy, Valerie Lecouturier, Sophie Ruiz, Catherine Berry, Timothy Tibbitts, Hanne Andersen, Anthony Cook, Alan Dodson, Laurent Pessaint, Alex Van Ry, Marguerite Koutsoukos, Cindy Gutzeit, I-Ting Teng, Tongqing Zhou, Dapeng Li, Barton F. Haynes, Peter D. Kwong, Adrian McDermott, Mark G. Lewis, Tong Ming Fu, Roman Chicz, Robbert van der Most, Kizzmekia S. Corbett, Mehul S. Suthar, Galit Alter, Mario Roederer, Nancy J. Sullivan, Daniel C. Douek, Barney S. Graham, Danilo Casimiro, Robert A. Seder

**Author notes:** Corresponding author Cellular Immunology Section, Vaccine Research Center, National Institute of Allergy and Infectious Disease, National Institutes of Health, 40 Convent Drive, MSC 3025, Building 40, Room 3512, Bethesda, MD 20892.

## Abstract

Adjuvanted soluble protein vaccines have been used extensively in humans for protection against various viral infections based on their robust induction of antibody responses. Here, soluble prefusion-stabilized spike trimers (preS dTM) from the severe acute respiratory syndrome coronavirus (SARS-CoV-2) were formulated with the adjuvant AS03 and administered twice to nonhuman primates (NHP). Binding and functional neutralization assays and systems serology revealed that NHP developed AS03-dependent multi-functional humoral responses that targeted multiple spike domains and bound to a variety of antibody F_C_ receptors mediating effector functions *in vitro*. Pseudovirus and live virus neutralizing IC_50_ titers were on average greater than 1000 and significantly higher than a panel of human convalescent sera. NHP were challenged intranasally and intratracheally with a high dose (3×10^6^ PFU) of SARS-CoV-2 (USA-WA1/2020 isolate). Two days post-challenge, vaccinated NHP showed rapid control of viral replication in both the upper and lower airways. Notably, vaccinated NHP also had increased spike-specific IgG antibody responses in the lung as early as 2 days post challenge. Moreover, vaccine-induced IgG mediated protection from SARS-CoV-2 challenge following passive transfer to hamsters. These data show that antibodies induced by the AS03-adjuvanted preS dTM vaccine are sufficient to mediate protection against SARS-CoV-2 and support the evaluation of this vaccine in human clinical trials.

## Introduction

The 2019 outbreak of coronavirus disease (COVID-19) caused by the novel severe acute respiratory syndrome coronavirus (SARS-CoV-2) has become a global pandemic with 112,849,164 infections and 2,503,390 deaths across 192 countries, as of February 25, 2021^1^. An effective prophylactic vaccine remains the most effective public health measure for controlling disease spread^2^. To that end, two mRNA vaccines^3, 4^ have received emergency use authorization from the FDA based on clinical efficacy of greater than 90% in the US. In addition, adenovirus-based vaccines have been approved for use in the EU, United Kingdom^5^ and Russia^6^, and an inactivated virus vaccine is approved in China^7^. Other candidates based on protein are currently in clinical testing^8^. With the exception of the inactivated virus vaccines^9, 10^, these approved and clinical-phase candidate vaccines use only the coronavirus spike (S) protein as their immunogen.

Spike is a surface membrane-bound trimer that, by electron microscopy, gives viral particles a characteristic halo from which its family name “corona” is derived^11^. It is a type-1 viral membrane fusion protein that exists in a metastable prefusion conformation and undergoes a dramatic structural rearrangement upon engagement of the receptor binding domain (RBD) with its receptor, angiotensin-converting enzyme 2 (ACE2)^12,13,14^, ultimately leading to membrane fusion. It has been shown that antibodies directed against the RBD can neutralize incoming virus by preventing receptor recognition and thus entry^15,16,17,18,19^. Because the RBD, as well as other regions such as the N-terminal domain (NTD) may contain neutralizing epitopes^20, 21^, the full-length spike is a preferred target antigen for vaccine development. Based upon successful structure-based immunogen designs for SARS-CoV and Middle Eastern Respiratory Virus (MERS) vaccines^22, 23^, mutations have been introduced to block cleavage of S into S1 and S2 subunits and stabilize a region between the central helix and heptad repeat 1, giving rise to homogeneous S protein trimers in the prefusion conformation^24^. This construct, referred to as S-2P, is the basis for several SARS-CoV-2 vaccine candidates being delivered by adenoviral vectors^25^, displayed on nanoparticles^26^, or encoded by mRNA^27,28,29^.

In contrast to vectored gene delivery vaccine platforms, adjuvanted soluble protein vaccine formulations have been approved for clinical use against several viral infections^30,31,32^ and have a long history of being used across all age groups. Soluble protein subunit vaccines will likely require a potent adjuvant to elicit strong T and B cell responses^33^. Here we have formulated a soluble S-2P-derived protein with the well-characterized adjuvant, AS03, an oil-in-water emulsion composed of squalene, polysorbate 80, and α-tocopherol. AS03 potently induces antibodies and has been shown to increase vaccine durability, promote heterologous strain cross-reactivity^34^, and to have dose-sparing effects^35,36,37,38,39^. It was licensed for use in vaccines against pandemic influenza in Europe, with approximately 90 million doses administered ^35, 39,40,41^. Therefore, in this study, AS03-adjuvanted soluble S-2P trimers were evaluated for NHP immunogenicity and protection following SARS-CoV-2 challenge in advance of clinical trials. To date several advanced vaccine candidates have been characterized for the magnitude, quality and efficacy of the immune responses they elicit^25, 28, 42, 43^. Here we performed a thorough characterization humoral and cellular responses in the upper and lower respiratory tracts following vaccination and challenge. These studies establish that vaccine-induced antibody is sufficient for protection and highlight its role in rapid control of lower airway viral replication.

## Results

### Soluble spike trimers are immunogenic when adjuvanted with AS03

To create a SARS-CoV-2 protein vaccine, the S-2P stabilizing mutations were used as previously described^24^; the trimer was then expressed as a soluble protein by replacing the transmembrane domain with a T4 foldon domain, which has been shown to assist in trimerization of type-1 membrane fusion proteins^44, 45^ (Fig. 1a). The resulting soluble trimeric protein immunogen is thus referred to as prefusion transmembrane-deleted spike, or preS dTM. PreS dTM trimers were then formulated by admixing with the oil-in-water emulsion, AS03. NHP were immunized intramuscularly three weeks apart with or without AS03 to confirm its role for improving antibody responses. AS03 was critical for the induction of high magnitude S-2P IgG binding and neutralization titers (Fig. S1a-b), and S-2P-specific IgA and IgG B cell responses (Fig. S1c-f). Systems serology was also performed to assess the quantitative and qualitative effector functions of vaccine responses induced by AS03. Antibodies bound to a broad array of human antibody F_C_ receptors and enabled F_C_-mediated effector functions such as phagocytosis and complement activation (Fig. S1g,h). AS03 strongly enhanced all F_C_ functions equally with no skewing to a particular receptor or function. Collectively these data establish the critical role of the AS03 adjuvant for improving the magnitude and quality of antibody responses.

**Figure 1.**
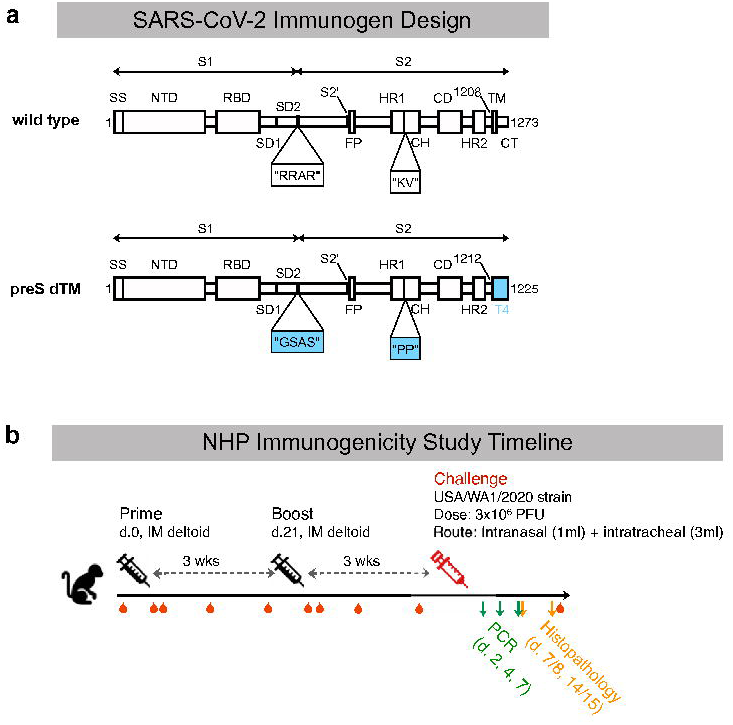
Vaccine design and study outline. **a**, Schematic of the SARS-CoV-2 spike protein, preS dTM, with stabilizing mutations at the S1/S2 furin cleavage site and the heptad repeat region; the transmembrane domain was replaced with a T4 trimerization domain. **b**, Schematic of NHP immunogenicity and challenge study. Immunizations were given at study weeks 0, 3; challenge was performed at study week 6; blood draws approximated by red droplets; PCR and necropsy/histopathology approximated by arrows.

To study protective efficacy and perform a wider assessment of immunogenicity, rhesus macaques were immunized with 4 or 12 µg AS03-adjuvanted preS dTM; PBS was administered as a negative control (Fig. 1b). Animals did not experience any abnormal body weight or temperature changes in response to vaccination (Supplementary Table 1). Serum binding titers were detectible two weeks after the first immunization at levels that approximated those found in human convalescent donor sera (HCS) from two different benchmark cohorts; endpoint binding titers were significantly increased from 2.9×10^3^ to 7.4×10^4^ following the second immunization in the high dose group (Fig. 2a). Notably there was no significant dose response between the 4 and 12 µg dose groups. In terms of the breadth of binding, antibody responses were observed to the S1 region and more specifically to the RBD and NTD (Fig. 2b).

**Figure 2.**
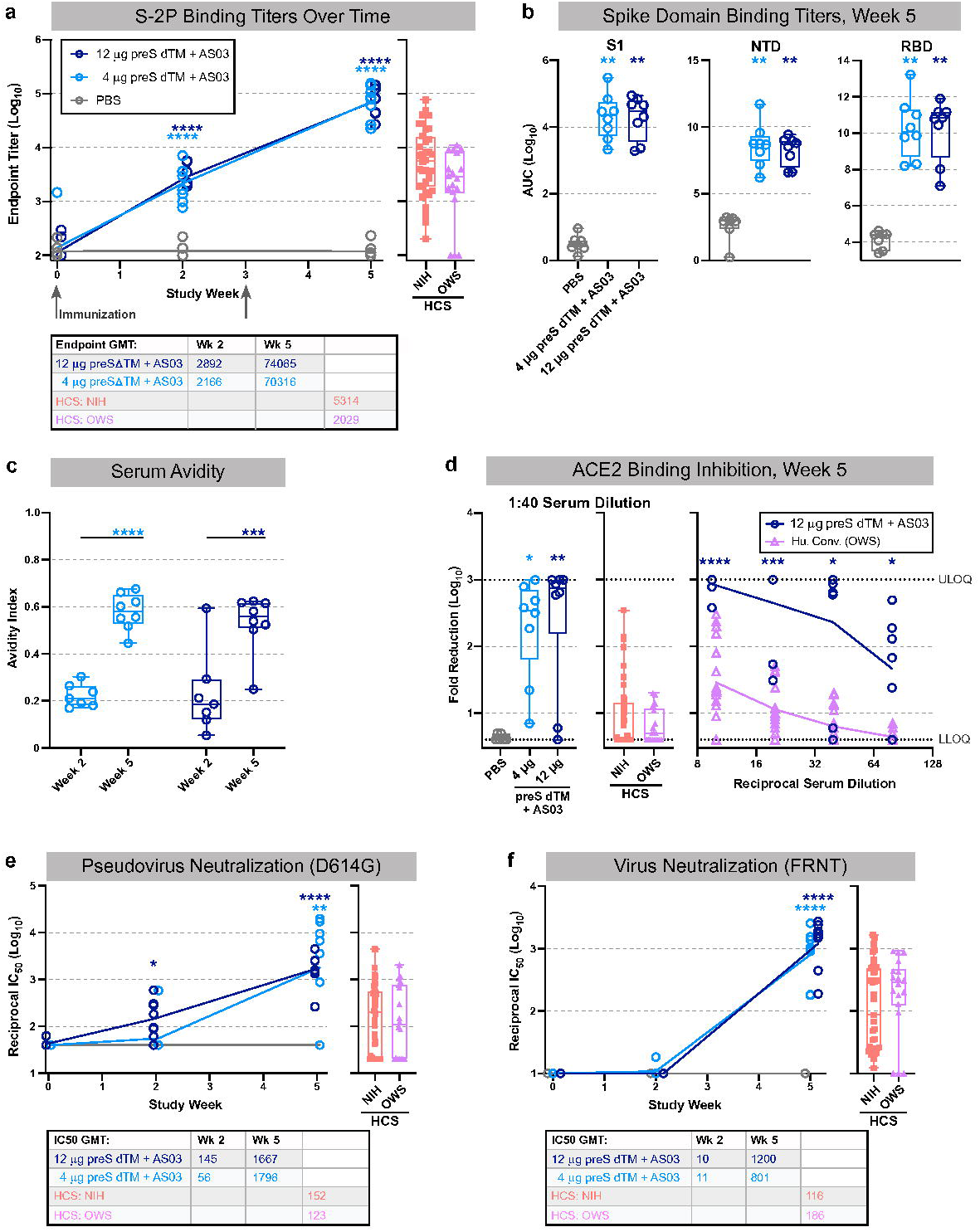
Serology following vaccination with AS03-adjuvanted preS dTM. Rhesus macaques were immunized with 4 or 12 μg of preS dTM adjuvanted with AS03 adjuvant at weeks 0 and 3. **a**, Endpoint binding titers following pre-vaccination (week 0) or post prime (week 2) and boost (week 5). **b,** Binding titers to the S1 domain, N-terminal domain (RBD) or receptor binding domain (RBD) at week 5. **c**, Avidity index at weeks 2 and 5. **d**, Plasma inhibition of ACE2 binding to spike; week 5 vaccine dose response at 1:40 dilution (left graph) or over multiple dilutions (right graph) are shown. Dotted lines indicate upper and lower limits of quantitation. **e**, Pseudovirus neutralization over time; IC_50_ values are plotted. **f**, Live virus neutralization over time; IC_50_ values are plotted. Symbols represent individual animals; box plots indicate the median and interquartile range; whiskers indicate minimum and maximum data points. Geometric mean values for binding and neutralization are indicated in tables below each graph. Two human convalescent serum (HCS) panels are plotted for comparison. Asterisks indicate significance compared to the PBS control group as follows: *, p<0.05; **, p<0.01; ***, p<0.001 ****, p<0.0001.

### Soluble spike trimers adjuvanted with AS03 induce neutralizing antibody responses

The next series of studies focused on functional antibody responses following AS03-adjuvanted preS dTM vaccination. First, the second immunization significantly improved serum avidity to S-2P in both dose groups (Fig 2c). Sera from both vaccine dose groups also showed ∼100-fold higher competition with ACE2 for binding to the RBD, compared to HCS (Fig. 2d). Inhibition of viral entry was next assessed using a pseudotyped reporter virus. While neutralization was low or undetectable in most animals after the first immunization, reciprocal titers over 10^3^ were achieved in nearly all animals following the boost (Fig. 2E). Similar results were seen with neutralization of live virus in a focus reduction neutralization titer assay, and these responses were generally 10-fold higher than those of HCS (Fig. 2F). Based on recent outbreaks of variant strains, we assessed neutralization against the B.1.1.7 “UK” and B.1.351 “South African” variants. Notably, there was ∼2-fold decrease against the B.1.1.7 variant, and ∼5 to 10 - fold reduction against the B.1.351 variant (Fig. S2).

### Soluble spike trimers adjuvanted with AS03 induce a mixed CD4 T cell response

Since adjuvants also have an important effect on the magnitude and quality of CD4 T cells, we measured the frequency of spike specific memory T_H_1 (IL-2, TNF, and IFN*γ*), T_H_2 (IL-4, IL-13), Th17 (IL-17) and T_FH_ (CXCR5^+^, PD-1^+^, ICOS^+^) IL-21 and CD40 ligand responses from PBMCs by multi-parameter flow cytometry. Two weeks following the boost (week 5), both T_H_1 and T_H_2 cytokines were detected (Fig. 3a). In assessing individual cytokines, the T_H_1 response was comprised mostly of IL-2 and TNF with minimal IFN*γ* production, indicative of a “T_H_0” phenotype^46, 47^ (Fig. 3b). Of note, antigen-specific IL-21 production and CD40 ligand expression were detected in both CD4 memory and T_FH_-gated PMBC subsets, supporting their role in the robust antibody responses induced following vaccination (Fig. 3c). To further analyze cytokine production on a single cell-basis, Boolean gating was used to show the various combinations of cytokines (Fig. 3D). Greater than ∼89% were CD40L+, a sensitive marker for antigen specific cells. Notably, just 6.5% of cells produced only T_H_2 cytokines, while ∼27% produced combinations of IL-2 or TNF, and IL-4 or IL-13, characterized as a mixed or “T_H_0” phenotype. CD8 T cell responses were largely undetectable (Fig. 3e).

**Figure 3.**
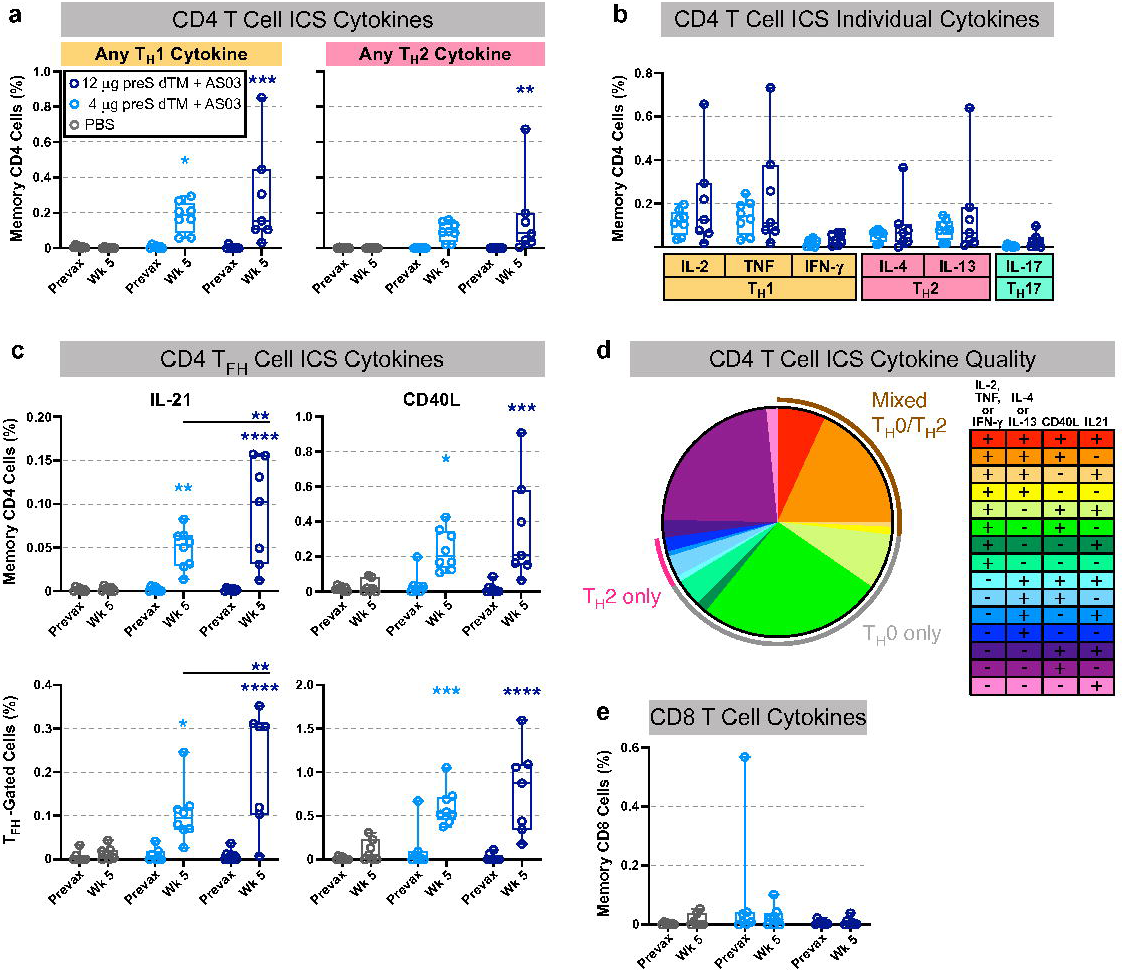
T cell responses following vaccination with AS03-adjuvanted preS dTM. T cell responses in rhesus macaques immunized with 4 or 12 ug of preS dTM adjuvanted with AS03 adjuvant at weeks 0 and 3. Cells taken before immunization (prevax) or at week 5 were stimulated with the S1 peptide pool covering the spike protein, then assessed by intracellular cytokine staining. **a**, Percent of memory CD4 T cells expressing any T_H_1 cytokine (IL-2, TNF, or IFN*γ*; left graph), or any T_H_2 cytokine (IL-4 or IL-13; right graph). **b**, Percent of memory CD4 T cells expressing the indicated cytokine. **c**, Percent of CD4 T cells expressing the T_FH_ markers IL-21 (left graphs) or CD40L (right graphs) in all memory CD4 cells (top row) or the T_FH_ subset (bottom row). **d**, Proportion of memory CD4 T cells expressing any T_H_1 (IL-2, TNF, or IFN*γ*), T_H_2 (IL-4 or IL-13), or T_FH_ (IL-21 or CD40L) markers by ICS Boolean gating; week 5 responses from both vaccine dose groups are averaged. Pie arcs indicate the proportion of cells expressing any T_H_1 and T_H_2 cytokines in the same cell (brown arc); T_H_1 cytokines only (grey arc), or T_H_2 cytokines only (pink arc). **e**, Percent of memory CD8 T cells expressing any T_H_1 cytokine (IL-2, TNF, or IFN*γ*). Symbols represent individual animals; box plots indicate the median and interquartile range; whiskers indicate minimum and maximum data points. Asterisks indicate significance compared to the PBS control group (unless otherwise indicated) as follows: *, p<0.05; **, p<0.01; ***, p<0.001 ****, p<0.0001.

### Soluble spike trimers adjuvanted with AS03 protect NHP from high dose SARS-CoV-2 challenge

Prior NHP vaccine studies^25, 48^ have used varying doses of the USA-WA1/2020 isolate ranging from 10^4^ to 10^6^ PFU for nasal and intratracheal challenge. In addition, passaging of the USA-WA1/2020 isolate has led to mutations in the furin cleavage site that can limit pathogenicity and results in variation of the amount and duration of infection in NHP. Thus, in this study, NHP were challenged 3 weeks following the boost with a new sequence-validated stock of the USA-WA1/2020 isolate that was administered at a high dose of 3×10^6^ PFU SARS-CoV-2 given intranasally and intratracheally. Lower airway protection was assessed using subgenomic RNA (sgRNA), as a quantitative metric of replicating virus^49^ in BAL (Fig. 4a). At day 2, 6/7 (86%) PBS control animals had detectable sgRNA, compared to 6/8 (75%) and 3/8 (38%) in the 4 µg and 12 µg dose groups, respectively. By day 4, 5/7 (71%) of PBS control animals were positive, but sgRNA was significantly reduced to 2/8 (25%) or 0/8 (0%) in the 4 µg and 12 µg vaccine dose groups, respectively. By day 7, sgRNA was detectable in 4/7 (57%) PBS controls but only 1/16 (6%) vaccinated animals. To assess upper airway protection, sgRNA was quantified in nasal swab extracts (Fig. 4b). Both vaccine groups showed significant sgRNA reduction of ∼1-3 Log_10_ on day 2. By day 4, 5/7 (71%) PBS controls had detectible sgRNA (10^4^), while only 2/8 (25%) and 0/8 (0%) vaccinated NHP had detectable sgRNA in the 4 µg and 12 µg dose groups, respectively. Thus, AS03-adjuvanted preS dTM provided significant protection in the upper and lower airways from this robust SARS-CoV-2 challenge.

**Figure 4.**
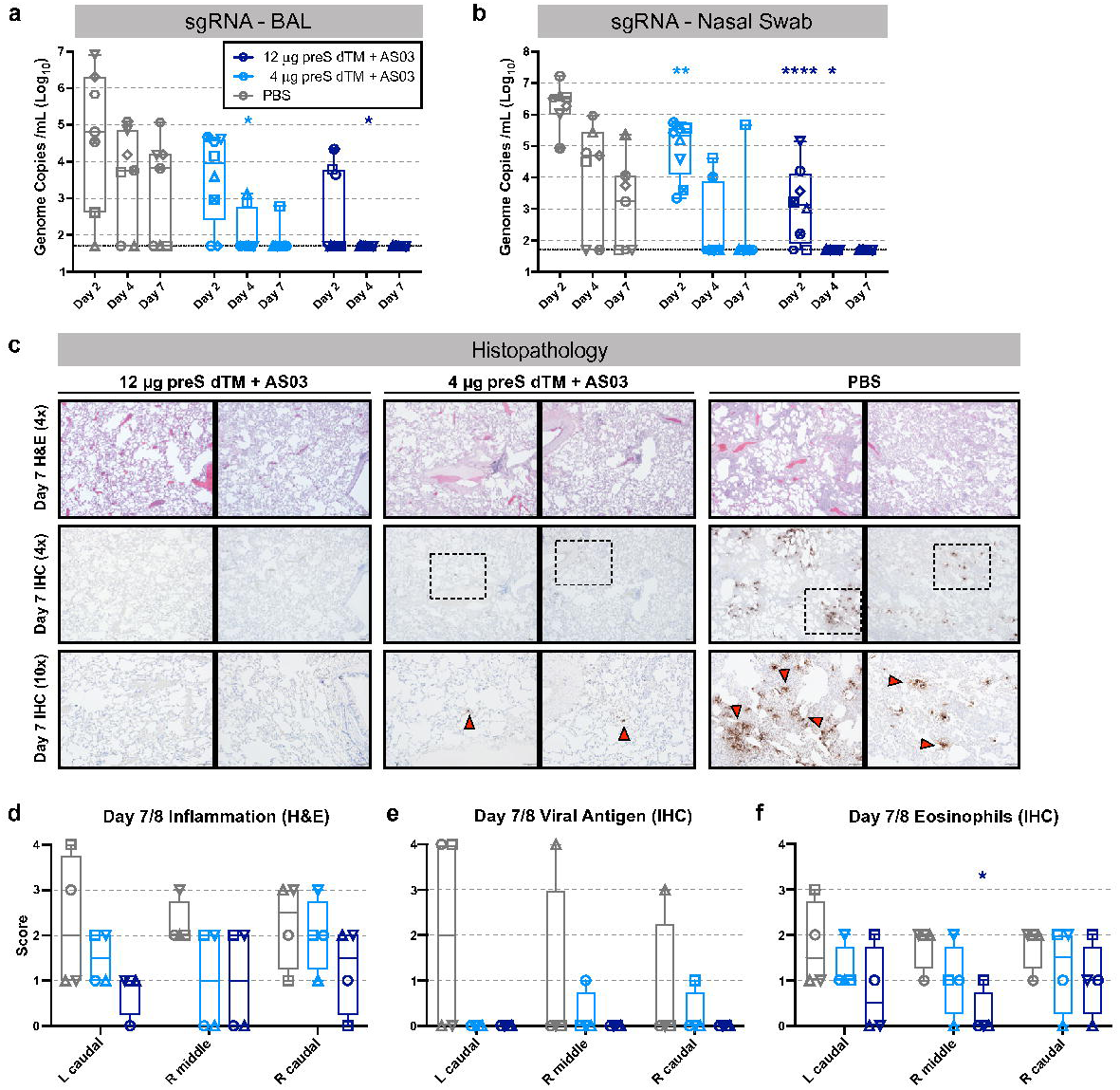
Vaccination with AS03-adjuvanted preS dTM protects NHP from SARS-CoV-2 challenge. Rhesus macaques immunized with 4 or 12 μg of AS03-adjuvanted preS dTM were challenged with 3×10^6^ PFU SARS-CoV-2 by the intranasal and intratracheal routes. SARS-CoV-2 subgenomic RNA (sgRNA) in BAL (**a**) and nasal swabs (**b**) at 2, 4, and 7 days following challenge. Symbols represent individual animals; bars indicate group geometric means. Dotted lines indicate lower limit of quantitation. (**C**) Histopathological analysis at 7 days post challenge. Representative images from lung sections from 2 animals per group analyzed by H&E stain for inflammation (top row, 4x magnification), or immunohistochemical (IHC) staining for viral antigen (middle row, 4x magnification; bottom row, 10x magnification). Red arrows indicate foci of viral antigen. **d-f**, Quantification of histopathology for 4 animals at days 7, 8 post challenge. Inflammation from H&E staining, **d**; viral antigen from IHC, **e**; and eosinophils from IHC, **f**. Symbols represent individual animals; box plots indicate the median and interquartile range; whiskers indicate minimum and maximum data points. Asterisks indicate significance compared to the PBS control group as follows: *, p<0.05; **, p<0.01; ****, p<0.0001.

To further substantiate the vaccine protection, lung tissue was analyzed for viral antigen, inflammation, and eosinophil infiltration in half of the animals in each group 7 days following challenge (Fig. 4c-e). Viral antigen was detected in at least 1 lobe of 3/4 PBS control animals, while antigen was undetectable in the high-dose vaccinated animals and had only limited detection in 2 of the low-dose vaccinated animals. Vaccination at either dose trended to reduce both tissue inflammation (Fig. 4d) and the presence of eosinophils (Fig. 4f), though these observations did not reach statistical significance.

### SARS-CoV-2 challenge boosts antibody titers in the lung

To further investigate how T cells or antibodies may have influenced protection in the respiratory tissues, we assessed T cell responses in the BAL and PBMC and antibody responses in the serum, BAL and nasal washes at various time points post challenge. Compared to the peak T cell responses after the second immunization at week 5 (restimulated with S peptides, Fig. 3a,b) memory PBMC CD4 and CD8 T cell responses were largely unchanged 7 to 14 days post challenge (Fig. S3a,b). However in BAL samples, spike-specific IL-2, IFN*γ*, and IL-13 recall responses were increased in the vaccinated groups compared to week 5, but not in the PBS controls (Fig. S3c,d). To assess the primary T cell response to infection, cells were restimulated with peptides to nucleoprotein, which is not present in the vaccine. Here we noted that the SARS-CoV-2 challenge induced a strong T_H_1 response in the BAL but not in PBMCs by day 14 that was specific to the PBS control animals (Fig. S3e,f). These data suggest that vaccine-elicited immune responses controlled the infection before a detectable primary T cell response could be generated in BAL or PBMCs.

Because vaccinated animals showed no detectable primary N-specific T cell response to the challenge in BAL or PBMCs, it suggested rapid control of infection by the vaccine in the airways, which we hypothesized might be mediated by antibodies. To assess this, we performed a kinetic analysis of antibody responses in BAL and nasal washes up to two weeks post challenge. Remarkably, S-2P IgG binding titers were significantly increased in the BAL from vaccinated animals just 2 days post challenge and remained higher than the control animals through day 7 (Fig. 5a). In contrast, IgA and IgG responses to the challenge developed only by day 14 in the PBS control animals consistent with the kinetics of a primary response (Fig. 5a,b). Notably this anamnestic response in the vaccinated animals was specific to the BAL, as there was no increase in S-2P IgG titers in the nasal wash (Fig. 5c) or the serum (Fig. 5d) post challenge. Importantly, the primary antibody response was evident in blood and upper and lower airways by day 14 in the PBS control animals. Based on the rapid anamnestic response in BAL in the vaccine groups, we next determined whether this was specific to S-2P antibodies, or if the challenge was causing a general increase in IgG. Indeed, there was an increase in total IgG in BAL of vaccinated animals at 2 days post challenge, continuing through day 4 and decreasing by day 7, while PBS control animals had a smaller increase on day 4 only (Fig. S4a). We next assessed whether other antibody specificities were increased in BAL post challenge. Because all NHP used in our studies have had earlier vaccination against measles, we assessed measles antibody titers in BAL. Consistent with increases in spike and total IgG titers in BAL, vaccinated animals similarly showed a significant increase in measles antibodies on days 2 and 4 compared to pre-challenge, while several PBS control animals also increased on day 4 (Fig. S4b). Finally, to assess whether this increase in total IgG could be due to increased general transudation into the lung from the serum, we assessed the serum protein albumin levels in BAL. Albumin levels in BAL were not significantly increased in the vaccinated or PBS control animals following SARS-CoV-2 challenge (Fig. S4c). These data show that SARS-CoV-2 challenge leads to a rapid and transient local increase in IgG that occurs earlier in vaccinated animals.

**Figure 5.**
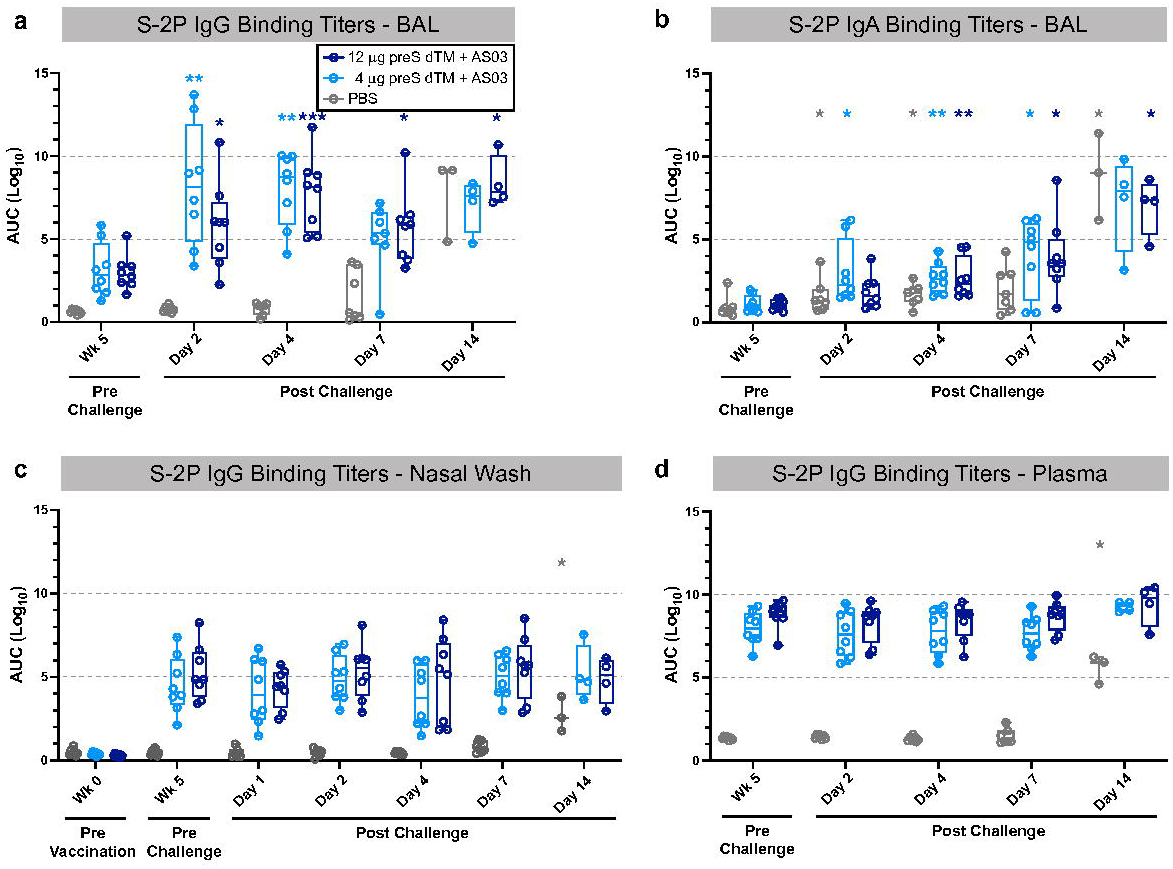
Anamnestic antibody responses in the lung following SARS-CoV-2 challenge. Bronchoalveolar lavage (BAL) supernatant was collected prior to challenge (week 5) and on days 2, 4, 7 and 14 following SARS-CoV-2 challenge. S-2P IgG (**a**) and IgA (**b**) binding titers in BAL samples. S-2P IgG binding titers in nasal washes (**c**) and plasma (**d**) taken pre- and post-challenge. Symbols represent individual animals; box plots indicate the median and interquartile range; whiskers indicate minimum and maximum data points. Asterisks indicate significance compared to the PBS control group as follows: *, p<0.05; **, p<0.01; ***, p<0.001.

### Vaccine induced IgG is sufficient to confer protection from SARS-CoV-2 challenge

Based on the high antibody and neutralizing titers in the blood and rapid anamnestic antibody responses in the BAL following challenge, we hypothesized that IgG was mediating protection. To directly assess whether vaccine induced antibody was sufficient to mediate protection, NHP IgG was purified from pooled plasma three weeks following the second vaccination just prior to the challenge and passively transferred to hamsters (Fig. 6a). A total of 10 mg or 2 mg total IgG per animal from AS03-adjvuanted preS dTM vaccinated NHP or from animals prior to vaccination as a negative control was administered to 8 individual hamsters per group. This resulted in approximately 125 mg and 25 mg IgG/kg bodyweight for the 10 and 2 mg dose groups, respectively (Fig. S5a). PBS was administered to an additional group as a negative control, and the highly potent and clinically approved SARS-CoV-2 mAb, LY-CoV555^50^ was administered to another group at 10 mg/kg as a positive control. Just prior to challenge, serum S-2P titers were confirmed in all but 2 animals, which were subsequently excluded (Fig. S5 b,c). Notably, the LY-CoV555 mAb recipient animals had higher ELISA binding titers than those that received polyclonal post-vaccination IgG. Hamsters that received only PBS prior to SARS-CoV-2 challenge lost an average of 10-15% of their bodyweight at day 6, a primary outcome measure of disease progression (Fig. 6b, Supplementary Table 2). Hamsters that received 10 mg of post-vaccination IgG had little weight loss and gained weight at a rate almost equivalent to that of the LY-CoV555 recipient hamsters. This protection was dose-dependent, as the animals that received 2 mg of post-vaccination IgG showed weight loss of ∼7% by day 6. Finally, individual animal serum S-2P binding titers were strongly correlated with body weight change, confirming the effect of IgG in protection from challenge (Fig. 6c). Taken together, these data show that the AS03-adjuvanted preS dTM vaccine elicited IgG sufficient to mediate protection *in vivo* against SARS-CoV-2 infection.

**Figure 6.**
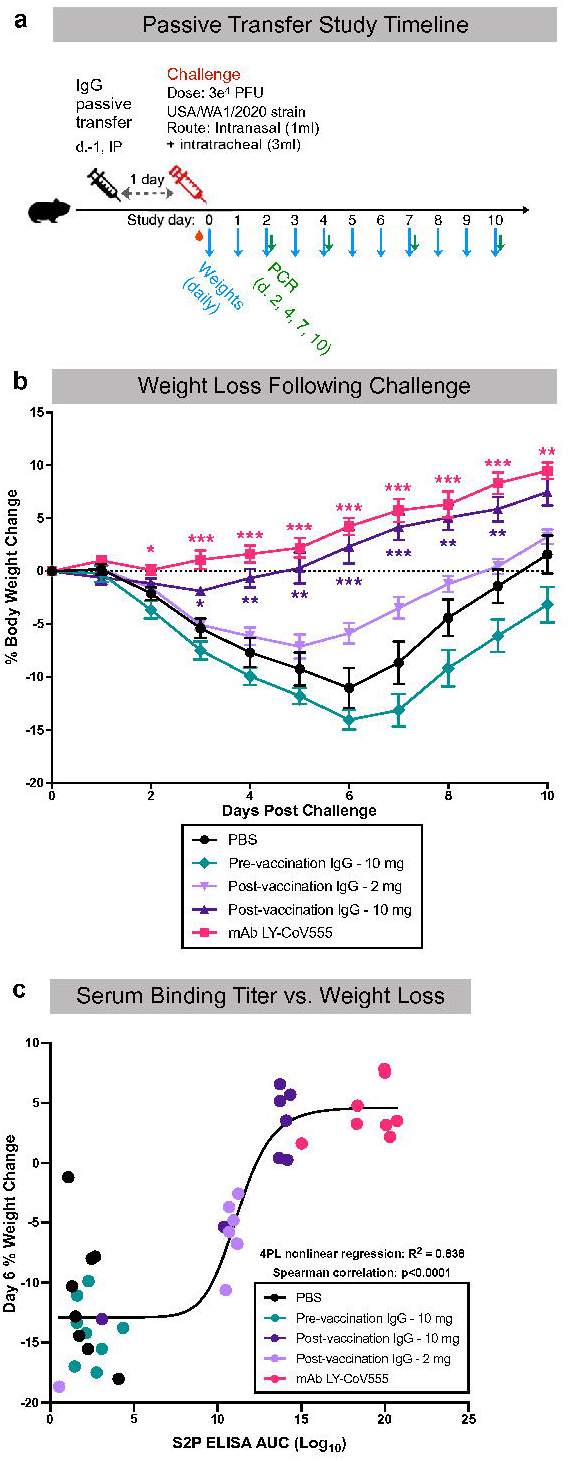
Passively transferred IgG from vaccinated NHP protects hamsters. Total IgG was isolated from pooled week 6 sera from rhesus macaques immunized with 3 μg of AS03-adjuvanted preS dTM. 10 or 2 mg total IgG was transferred to hamsters; 10 mg IgG from before (pre-) vaccination or PBS was transferred as a negative control for protection; 10 mg/kg mAb 555 was transferred as a positive control. Animals were then challenged 1 day later with SARS-CoV-2; body weight was recorded daily and oral swabs were taken for PCR on days 2, 4 and 7. **a**, Passive transfer study timeline. **b**, Daily change in body weight following challenge. Lines depict group mean body weight change from day 0; error bars represent SEM. **c**, Correlation between serum S-2P binding titer and percent weight loss on day 6 following SARS-CoV-2 challenge. Curve depicts a 4-parameter logistic fit of the data. Symbols represent individual animals; box plots indicate the median and interquartile range; whiskers indicate minimum and maximum data points. Asterisks indicate significance compared to the PBS control group at each time point: *, p<0.05; **, p<0.01; ***, p<0.001.

## Discussion

In this study, an AS03-adjuvanted soluble prefusion S protein vaccine formulation produced by Sanofi Pasteur and GlaxoSmithKline was evaluated for its ability to protect nonhuman primates against SARS-CoV-2 challenge in advance of clinical trials. Although SARS-CoV-2 mRNA- and adenovirus-based vaccine candidates have been authorized for emergency use in various countries, adjuvanted protein vaccines provide an additional vaccine platform to prevent disease that could be broadly useful in all age groups based on their long history and safety record with other viral infections. A key aspect of these studies was to provide new insights into the mechanisms of protection, most notably in the lung, which is critical for understanding how vaccines limit disease, a primary endpoint in all clinical trials.

By comparing the immune response to preS dTM formulations with and without AS03, it is clear that AS03 is critical for the induction of protective antibody responses, as has been previously observed with influenza^35, 36, 51, 52^ and respiratory syncytial virus (RSV)^53^. The CD4 responses to spike were primarily T_H_0 given the relative limited IFNγ production^46^ and T_H_2. In mouse studies, IL-4 and IL-13 production was also observed following vaccination with an inactivated influenza/AS03 formuation^54^. Previous human studies of AS03 with hepatitis B surface antigen^55^ and influenza hemagglutinin^56^ have observed strong IL-2 and TNF production with lower IFNγ responses. However, IFNγ responses were recently documented in humans with AS03 and a similar antigen (SCB-2019)^57^, suggesting that the CD4 profile might differ depending on the species and the antigen. Based on mouse and other animal models, vaccine-induced T_H_2 responses have been proposed to contribute to enhanced respiratory disease (ERD)^58,59,60^ as was observed in children given inactivated measles^61^ and RSV^62^ vaccines. Similarly, SARS vaccines formulated with the T_H_2-skewing adjuvant, alum, have been reported to induce immunopathology following challenge in mice, including eosinophilia^63^. Other studies suggest ERD is driven by nonfunctional and poorly matured antibodies^58, 64, 65^. Regardless of the mechanism, and in contrast to those findings, following SARS-CoV-2 challenge there was limited evidence of viral infection and there was a trend toward lower inflammation and eosinophil infiltration in lung tissue from vaccinated animals compared to PBS controls, indicating no enhanced disease.

A major focus of this study was to fully characterize the magnitude, quality and location of antibody responses. The NHP model has been used extensively for COVID-19 vaccine development and shares characteristics of mild human disease. Thus, a major advantage of using NHP is the ability to analyze immune responses in the mucosa of the upper and lower airways. Regarding the magnitude of antibody responses, pseudo- and live virus-neutralization titers were above 10^3^ in most animals 2 weeks after the second immunization of AS03-adjuvanted preS dTM. These responses are comparable to prior studies using similar assays following 100 µg of the mRNA nanoparticle vaccines, mRNA1273^28^ and BNT162b2^48^ and importantly are superior to those from convalescent humans.

The AS03-adjvuanted preS dTM was able to rapidly reduce viral replication in both the upper and lower airways (BAL) by day 2, with no detectable virus in any of the animals in the high dose group by day 4. Comparing these results to other vaccines is difficult based on differences in the challenge stock dose and virulence. Here we have used a high challenge dose of 3×10^6^ PFU, which is 5-200 fold higher than the doses used to evaluate mRNA1273^28^, Ad26.COV2-S^25^, ChAdOx1 nCoV-19^43^, and NVX-CoV2373^42^, but similar to that used for BNT162b2^48^. The relatively higher challenge dose used here may in part explain why there was not complete reduction in viral titers at day 2, as was observed with other vaccines tested in NHP using lower challenge doses. Similarly we observed nasal swab titers of ∼2×10^6^ sgRNA copies/mL in the control animals at day 2, 10-50 fold higher than previously reported in other NHP studies by us and others^28, 43^, likely reflecting both the more virulent challenge and improved sgRNA extraction methods.

Because significant protection was conferred by vaccination with AS03-adjuvanted preS dTM, it was critical to investigate the potential mechanisms for this effect. Consistent with data from most protein subunit vaccine studies in humans with AS03 and other adjuvants, CD8 T cell responses were undetectable. These data suggest they would have a limited role in primary, rapid protection in this model. By contrast, potent antigen-specific CD4 T cells were comprised of Th0, T_H_2 and T_FH_ cytokines and expressed CD40L, which are optimal for generating robust antibody responses. In terms of having effector functions to mediate protection, a direct protective role of CD4 T cells would have required cell depletion prior to challenge and was not assessed in this study.

Based on the early control of infection seen by day 2, it is likely that antibodies had a significant role in neutralizing replicating virus. That neutralizing antibodies could protect against challenge has been shown in correlative analyses with other SARS-CoV-2 vaccine candidates^28, 66^. Moreover, prior studies have shown a protective effect from passively transferred convalescent sera to hamsters^67^ and NHP^68^. However, here is the first demonstration that vaccine-elicited IgG could protect against challenge, providing direct evidence that antibodies are sufficient to mediate protection. An additional finding that substantiates a role for antibodies was the rapid increase in IgG responses in the lung as early as 2 days following challenge in NHP. Mucosal antibody responses to vaccination have been well documented, including in response to polio^69,70,71^, influenza^72, 73^ and RSV^74, 75^ vaccines. Moreover, neutralizing BAL responses have been observed in humans in the initial days after symptom onset^76^. However, to the best of our knowledge, this is the first report of vaccine-directed anamnestic lung responses to challenge. This rapid increase in spike IgG appears specific to the BAL compartment and was not observed in the upper airway or serum post-challenge. A notable finding was that there was also a transient increase in total IgG titers and measles antibody from prior vaccination in the BAL at day 2, yet albumin levels were not increased. These data suggest that the increase in antibodies may not be due to a general transudation of proteins from increased vascular leakage or bulk vesicular transport. We speculate that this increase in total and antigen-specific IgG could occur through a general activation of memory B cells in the lung, perhaps directly from TLR-sensing of viral RNA^77^ or through bystander activation from activated S-specific CD4 T cells. Together this could explain why IgG titers were higher in vaccinated animals on day 2, while PBS controls showed a smaller increase in total and measles IgG only on day 4. While antibody secreting cell (ASC) enumeration was not possible in this study, future studies will examine changes in lung resident ASCs and investigate whether an increase in BAL antibodies is specific to this vaccine or is generalizable to other vaccine platforms. Nevertheless, these data highlight the potential role of ASCs in contributing to control of viral infections in the lung.

In conclusion, this report highlights that potent serum antibody responses are induced by soluble S trimers formulated with the oil-in-water emulsion adjuvant AS03, which conferred protection in the upper and lower airways following SARS-CoV-2 challenge. This vaccine induced IgG responses sufficient to protect from SARS-CoV-2 challenge, and the rapid anamnestic response in the lower airway likely contributed to protection at this site. These data support the clinical development of the AS03-adjuvanted preS dTM vaccine for limiting SARS-CoV-2 infection and protecting against COVID-19 morbidity and mortality.

## Materials and Methods

### Immunogen

The SARS-CoV-2 recombinant vaccine candidate consists of purified recombinant prefusion spike (S) protein (SARS-CoV-2 prefusion-stabilized S delta TM (preS dTM)) adjuvanted with AS03. The preS dTM was produced from a Sanofi Pasteur proprietary cell culture technology based on the insect cell – baculovirus system, referred to as the Baculovirus Expression Vector System (BEVS). The preS dTM sequence was designed based on the Wuhan YP_009724390.1 strain S sequence, but modified to improve the conformation, stability, trimerization and facilitate the purification. The modifications comprise mutation of the S1/S2 furin cleavage site, introduction of two proline mutations in the C-terminal region of S2 domain, deletion of the transmembrane and cytoplasmic region and replacement by the T4-foldon trimerization domain^24^. Briefly, the modified sequence was cloned into a baculovirus transfer plasmid, which was then used to generate a recombinant baculovirus containing the gene of interest. The recombinant baculovirus was first amplified in expresSF+ insect cells prior to infecting a large scale expresSF+ insect cells culture in suspension. After incubation, the recombinant protein was purified from the supernatant using several affinity and chromatography columns. Based on an ACE2 binding assay, the preS dTM used in the study were quantified at 4 µg and 12 µg for a total protein content of 5 and 15 µg, for the low and high doses respectively.

### Adjuvant and formulation

AS03 is an Adjuvant System composed of *α*-tocopherol, squalene and polysorbate 80 in an oil-in-water emulsion^78^. Vaccine doses were formulated by diluting the appropriate dose of preS dTM with PBS-tween to 250 µL, then mixing with 250 µL AS03, followed by inversion five times for a final volume of 500 µL. Each dose of AS03 contains 11.86 mg *α*-tocopherol, 10.69 mg squalene and 4.86 mg polysorbate-80 (Tween 80) in PBS.

### Animals, immunizations, challenges and sampling

Rhesus macaques were randomized into groups of 8 based on age and body weight; each group had 2 females and 6 males, except for the PBS control group, which only had 5 males. All animals had a history of measles vaccination. SARS-CoV-2 vaccine formulations were administered by intramuscular injection into the right deltoid for both immunizations, three weeks apart. Whole blood was collected weekly into EDTA-containing tubes and also 1, 4 and 7 days after each immunization for complete blood count/chemistry profiling. PBMC and plasma were then collected from whole blood following ficoll purification. For SARS-CoV-2 challenge, virus was obtained from Operation Warp Speed: strain-2019-nCoV/USA-WA1/2020, Lot# 70038893; BEI resources catalog no. NR-53780. Virus was inoculated intranasally (0.5 mL per nostril) and intratracheally (3 mL) for a total of 3×10^6^ PFU per animal. Bronchiolar/alveolar lavage (BAL) sampling was performed 5 weeks after vaccination and 2, 4 and 7 days after challenge. Nasal swabs for sgRNA PCR were taken 5 weeks after vaccination and 2, 4 and 7 days after challenge; nasal washes (∼5 mL PBS) were collected at weeks 0 and 5, and 1, 2, 4, 7 and 14 days post challenge. Half the animals in each group were necropsied at day 7 and 14 post challenge, where lung tissue was collected for histopathology.

In a separate study, 4 groups of 6 rhesus macaques were similarly vaccinated, but with lower doses of 1.3 and 3.9 µg preS dTM + AS03, or 3.9 µg preS dTM without AS03, or PBS alone. Animals were then boosted at week 3 with 2 and 6.1 µg preS dTM + AS03, or 6.1 µg preS dTM without AS03, or PBS alone. Immunogenicity data from this study are found in Figure S2. IgG were purified from animals given 3 µg preS dTM + AS03 either before immunization or 3 weeks following the second immunization for use in passive transfer to hamsters, Figure 6.

For passive transfer studies, 6–8-week-old Golden Syrian hamsters were randomized into groups of 8 based on weight, each group having 4 males and 4 females. IgG was passively transferred by intraperitoneal injection 1 day prior to challenge. SARS-CoV-2 challenge virus (strain-2019-nCoV/USA-WA1/2020, Lot# 70038893; BEI resources catalog no. NR-53780) was introduced intranasally at a dose of 3×10^4^ PFU administered in a final volume of 100 µL and split between each nostril. Body weight and clinical observations were made daily; serum was sampled just prior to challenge; oral swabs were taken 0, 2, 4, 7 and 10 days post challenge for viral load determination.

### Ethics statement

Macaques were housed at the NIH (for immunizations) and Bioqual, Inc. (for challenge); hamsters were housed at Bioqual, Inc. All animals were cared for in accordance with American Association for Accreditation of Laboratory Animal Care standards in accredited facilities. All animal procedures were performed according to protocols approved by the Institutional Animal Care and Use Committees of the National Institute of Allergy and Infectious Diseases, National Institutes of Health and Bioqual, Inc. NHP studies were performed under NIH animal study protocol #VRC-20-870; hamster studies were performed under animal study protocol #VRC-20-872.

### Human convalescent sera

Two panels of samples from human patients who had recovered from SARS-CoV-2 disease were used in parallel. The first panel referred to as “NIH” has been described previously^28^. Additionally, an 18-sample panel collected by Operation Warp Speed and distributed by Battelle and BEI Resources was also used, referred to here as “OWS”. Informed consent was obtained from all participants. Participants had a history of laboratory-confirmed SARS-CoV-2 infection before they provided serum.

### Serology

Antibody titers to various SARS-CoV-2 derived antigens were assayed as previously described^28^. Briefly, endpoint binding titers were measured by standard sandwich ELISA using proline-stabilized spike protein (S-2P). Binding to SARS-CoV-2 S1, receptor-binding domain (RBD), and N-terminal domain (NTD) spike subdomains was performed using Mesoscale Discovery ELISA^28^, using biotinylated subdomain proteins prepared as described previously^79^.

### Total IgG antibody titers were quantitated by using the Human IgG ELISA ^BASIC^ kit

(ALP) (Mabtech) following manufactures directions. Samples were read by MSD plate reader (Sector Imager 600). Antibody titers to measles were quantitated by using Monkey Anti-Measles IgG ELISA kit (Alpha Diagnostics International) following manufactures directions. The optical density (OD) of each well was read at 450 nm by SpectraMax Paradigm Multi-Mode Microplate Reader (Molecular Devices). Albumin levels were measured by bead-based single-plex assay using Luminex. The albumin analyte was selected and measured using MILLIPLEX MAP Human Kidney Injury Magnetic Bead Panel 2 (Millipore Sigma) following manufacturer’s instructions. Fluorescence data were collected on MAGPIX with Bio-Plex ManagerTM MP software (BioRad).

ACE2 binding inhibition was also performed via Mesoscale Discovery 384-well, 4-Spot Custom Serology SECTOR plates precoated with RBD; plasma was applied at a starting dilution of 1:10 followed by 10-fold serial dilutions. Binding was detected with SULFO-TAG–labeled ACE2 (Meso Scale Diagnostics).

For avidity analyses, plasma samples were heated at 56 °C for 45 min to complement-inactivate and reduce potential risk from any residual virus and immediately used or stored at -80°C for later use. 96-well plates (Nunc MaxisorpTM, Thermo Fisher) were coated with 100 µL of 1 µg/mL SARS-CoV-2 S-2P in 1x PBS for 16 hrs at 4°C. Plates were washed 3x in washing buffer (1x PBS, 0.2% Tween20, pH 7.4) using a Biotek 405 Microplate Washer and blocked with 200 µL blocking buffer (1 x PBS-T: 0.14 M NaCl, 0.0027 M KCl, 0.05% Tween 20, 0.010 M PO43-, pH 7.4) for 2 hrs at RT and washed 3x. Plasma/serum samples were serially diluted (starting dilution 1:100 and 4-fold dilutions) in blocking buffer and 100 µL was transferred to the plates. After 1 hour of incubation, plates were washed, and half of the samples were then incubated with 100 µL 1x PBS while the other half of the paired samples were treated with 100 µL 1.0 M sodium thiocyanate solution (NaSCN, Sigma-Aldrich) for 15 minutes at room temperature (RT) and washed 6x. Plates were incubated for 1 hr with 100 µL of goat-anti-human IgG (H+L, Cat#PA1-8463) or goat-anti-monkey IgG (H+L, Cat#A18811) secondary antibody conjugated to horseradish peroxidase (HRP, Thermo Fisher) in blocking buffer at 1:10,000 or 1:4,000 dilution, respectively. Plates were washed 3x and developed by addition of 100 µL KPL SureBlueTM TMB Microwell Peroxidase Substrate (1-Component, SeraCare, Cat#52-00-01) for 10 min. The reaction was quenched by addition of 100 µL 1 N H2SO4 and absorbance was measured at a test wavelength of 450 nm and reference wavelength of 650 nm using SoftMax Pro software version 6.5 on a Spectramax Paradigm Microplate reader (Molecular Devices). The avidity index (AI) was calculated using the ratio of the NaSCN-treated serum dilution giving an OD of 0.5 to the PBS-treated serum dilution giving an OD of 0.5 after 5PL curve-fitting in Graphad Prism. Reported AI is the average of two independent experiments, each containing duplicate samples. Samples with an OD < 0.5 could not be interpolated and were excluded from analysis.

To produce SARS-CoV-2 pseudotyped lentivirus, a codon-optimized CMV/R-SARS-CoV-2 S (Wuhan-1, GenBank: MN908947.3) plasmid was constructed and subsequently modified via site-directed mutagenesis to contain the D614G mutation. Further mutations were integrated into the D614G background to recapitulate the spike mutations of both the B.1.1.7 (UK) and B.1.351 (RSA) variants. Pseudoviruses were produced by co-transfecting HEK293T/17 cells (ATCC CRL-11268) with plasmids encoding a luciferase reporter, a lentivirus backbone, and the SARS-CoV-2 S genes into HEK293T/17 cells (ATCC CRL-11268) as previously described^80^. A human transmembrane protease serine 2 (TMPRSS2) plasmid was co-transfected to produce pseudovirus^81^. Neutralizing antibody responses in sera were assessed by pseudoneutralization assay as previously described^3, 22^. Briefly, heat-inactivated sera were serially diluted in duplicate, mixed with pseudovirus previously titrated to yield 10^4^ RLU, and incubated at 37°C and 5% CO2 for roughly 45 minutes. 293T-hACE2.mF cells were diluted to a concentration of 7.5 x 10^4^ cells/mL in DMEM (Gibco) supplemented with 10% Fetal Bovine Serum (FBS) and 1% Penicillin/Streptomycin and added to the sera-pseudovirus mixture. Seventy-two hours later, cells were lysed and luciferase activity (in relative light units (RLU)) was measured using a SpectraMaxL (Molecular Devices) luminometer. Percent neutralization was normalized, considering uninfected cells as 100% neutralization and cells infected with pseudovirus alone as 0% neutralization. ID_50_ titers were determined using a log(agonist) vs. normalized-response (variable slope) nonlinear regression model in Prism v8 (GraphPad).

For neutralization of authentic SARS-CoV-2 virus, a focus reduction neutralization titer (FRNT) assay was performed as previously described^82^. Plasma/serum were serially diluted (three-fold) in serum-free Dulbecco’s modified Eagle’s medium (DMEM) in duplicate wells and incubated with 100–200 FFU infectious clone derived SARS-CoV-2-mNG virus^83^ at 37 °C for 1 h. The antibody-virus mixture was added to VeroE6 cell (C1008, ATCC, #CRL-1586) monolayers seeded in 96-well blackout plates and incubated at 37 °C for 1 h. Post-incubation, the inoculum was removed and replaced with pre-warmed complete DMEM containing 0.85% methylcellulose. Plates were incubated at 37 °C for 24 h. After 24 h, methylcellulose overlay was removed, cells were washed twice with PBS and fixed with 2% paraformaldehyde in PBS for 30 min. at room temperature. Following fixation, plates were washed twice with PBS and foci were visualized on a fluorescence ELISPOT reader (CTL ImmunoSpot S6 Universal Analyzer) and enumerated using Viridot^84^. The neutralization titers were calculated as follows: 1 - (ratio of the mean number of foci in the presence of sera and foci at the highest dilution of respective sera sample). Each specimen was tested in two independent assays performed at different times. The FRNT-mNG_50_ titers were interpolated using a 4-parameter nonlinear regression in GraphPad Prism 8.4.3. Samples with an FRNT-mNG_50_ value that was below the limit of detection were plotted at 10. For these samples, this value was used in fold reduction calculations.

### Quantification of antigen-specific B cells

Cryopreserved PBMC were stained for antigen-specific B cells and subsets (Figure S6) using the following panel: IgM BUV395 (clone G20-127, Becton Dickenson), CD8 BUV665 (clone RPAT8, BD), CD56 BUV737 (clone NCAM16, BD), IgD FITC (Southern Biotech), IgA Dy405 (polyclonal, Jackson Immunoresearch), aqua LIVE/DEAD (Invitrogen), CD14 BV785 (clone M5E2, BioLegend), CD20 Alexa700-PE (clone 2H7, vaccine research center), IgG Alexa700 (clone G18-145, BD), CD3 Cy7APC (clone SP34-2, BD Pharmingen). Biotinylated prefusion-stabilized spike (S-2P) and spike subdomain (NTD, RBD) probes were produced as previously described^79^ and conjugated to streptavidin-labeled dyes (BD) to yield the following and streptavidin-conjugated B cell probes, NTD SA-BB700 (BD), RBD SA-BV650 (BD), and S-2P SA-APC (BD). PBMC were thawed into cRPMI + 10% FBS^85^, washed with PBS, and stained with aqua LIVE/DEAD kit in PBS for 20 min. at 4 °C. Staining was then completed with the remainder of the antibody and probe cocktail described above for 45 min. at 4 °C, and washed twice with PBS before flow cytometry.

### Intracellular Cytokine Staining

To measure vaccine-specific T cell responses, cryopreserved PBMC were thawed and rested overnight in a 37C/5% CO2 incubator. The next morning, cells were stimulated with SARS-CoV-2 spike protein (S1 peptide pools, JPT Peptide Technologies, Inc.) at a final concentration of 2 μg/ml in the presence of monensin and costimulatory antibodies anti-CD28 and -49d (clones CD28.2 and 9F10, BD Biosciences) for 6 hours. Negative controls received an equal concentration of DMSO (instead of peptides) and co-stimulatory antibodies. Intracellular cytokine staining and gating for CD4, CD8 was performed as previously described^86^ except the following monoclonal antibodies were added: PD-1 BUV737 (clone EH12.1, BD Biosciences) in place of PD-1 BV785, TNF-FITC (clone Mab11, BD Biosciences) in place of IL-5 BB515, and CD154 (CD40L) BV785 (clone 24-31, BioLegend). T_FH_ subsets were gated as CXCR5^+^, PD-1^+^, ICOS^+^. Aqua LIVE/DEAD kit (Invitrogen) was used to exclude dead cells. All antibodies were previously titrated to determine the optimal concentration.

### Flow cytometry

All phenotyping and ICS data were acquired on an BD FACSymphony flow cytometer and analyzed using FlowJo version 9.9.6 (Treestar, Inc., Ashland, OR).

### Luminex Isotype and FcR Binding Assay

To determine relative concentrations of antigen-specific antibody isotypes and Fc-receptor binding activity in the rhesus samples, a customized Luminex isotype assay was performed as previously described^87^. Antigens including SARS-CoV-2 full length spike (Eric Fischer, DFCI) and RBD (kindly provided by Aaron Schmidt, Ragon Institute) were covalently coupled to Luminex microplex carboxylated bead regions (Luminex Corporation) using NHS-ester linkages with Sulfo-NHS and EDC (Thermo Fisher Scientific) according to manufacturer recommendations. Immune complexes were formed by incubating antigen-coupled beads with diluted samples while rotating overnight. Mouse-anti-rhesus antibody detectors were then added for each antibody isotype (IgG1, IgG2, IgG3, IgG4, IgA, NIH Nonhuman Primate Reagent Resource supported by AI126683 and OD010976). Then, tertiary anti-mouse-IgG detector antibodies conjugated to PE were added. Flow cytometry was performed using a an iQue Plus Screener (Intellicyt) with a robot arm (PAA). Analysis of the flow cytometry data was performed using iQue Intellicyt software.

### Systems serology

In order to quantify antibody functionality of plasma samples, bead-based assays were used to measure antibody-dependent cellular phagocytosis (ADCP), antibody-dependent neutrophil phagocytosis (ADNP) and antibody-dependent complement deposition (ADCD), as previously described^88,89,90^. SARS-CoV-2 spike protein (kindly provided by Eric Fischer, DFCI) was coupled to fluorescent streptavidin beads (Thermo Fisher) and incubated with diluted plasma samples to allow antibody binding to occur. For ADCP, cultured human monocytes (THP-1 cell line, ATCC) were incubated with immune complexes to induce phagocytosis. For ADNP, primary PMBCs were isolated from whole blood from healthy donors using an ammonium-chloride-potassium (ACK) lysis buffer. After phagocytosis of immune complexes, neutrophils were stained with an anti-CD66b Pacific Blue detection antibody (Biolegend). For detection of complement deposition, lyophilized guinea pig complement (Cedarlane) was reconstituted according to manufacturer’s instructions and diluted in a gelatin veronal buffer with calcium and magnesium (Boston BioProducts). After antibody-dependent complement deposition occurred, C3 bound to immune complexes was detected with FITC-Conjugated Goat IgG Fraction to Guinea Pig Complement C3 (MP Biomedicals).

For quantification of antibody-dependent NK cell activation^91^, diluted plasma samples were incubated in Nunc MaxiSorp plates (Thermo Fisher Scientific) coated with antigen. Human NK cells were isolated the evening before using RosetteSep Human NK cell Enrichment cocktail (STEMCELL Technologies) from healthy buffy coat donors and incubated overnight with human recombinant Interleukin 15 (IL-15) (STEMCELL Technologies). NK cells were incubated with immune complexes, CD107a PE-Cy5 (BD), Golgi stop (BD) and Brefeldin A (BFA, Sigma-Aldrich). After incubation, cells were stained using anti-CD16 APC-Cy7 (BD), anti-CD56 PE-Cy7 (BD) and anti-CD3 Pacific Blue (BD), and then fixed (Perm A, Life Tech). Intracellular staining occurred with anti-IFNγ FITC (BD) and anti-MIP-1β PE (BD) after permeabilizing the NK cells with Perm B (Thermo Fisher). Flow cytometry acquisition of all assays was performed using the iQue Screener Plus (Intellicyt) and an S-LAB robot (PAA). For ADCP, phagocytosis events were gated on bead-positive cells. For ADNP, neutrophils were identified by gating on CD66b+ cells, phagocytosis was identified by gating on bead-positive cells. A phagocytosis score for ADCP and ADNP was calculated as (percentage of bead-positive cells) x (MFI of bead-positive cells) divided by 10,000. ADCD quantification was reported as MFI of FITC-anti-C3. For antibody-dependent NK activation, NK cells were identified by gating on CD3-, CD16+ and CD56+ cells. Data were reported as the percentage of cells positive for CD107a, IFNγ, and MIP-1β.

### Quantification of subgenomic RNA following challenge

Nasal swabs and bronchoalveolar-lavage (BAL) fluid were collected 2, 4, and 7 days post-challenge. At the time of collection, nasal swabs were frozen in 1mL of PBS containing 1 μL of SUPERase-In RNase Inhibitor (Invitrogen) and frozen at -80 °C until extraction. Nasal specimens were thawed at 55 °C, and the swab removed. The remaining PBS was mixed with 2 mL of RNAzol BD (Molecular Research Center) and 20 μL acetic acid. At the time of collection, 1 mL of BAL fluid was mixed with 1 mL of RNAzol BD containing 10 μL acetic acid and frozen at -80 °C until extraction. BAL specimens were thawed at room temperature and mixed with an additional 1 mL of RNAzol BD containing 10 μL acetic acid.

Total RNA was extracted from nasal specimens and BAL fluid using RNAzol BD Column Kits (Molecular Research Center) and eluted in 65 μL H_2_0. Subgenomic SARS-CoV-2 E (envelope) mRNA was quantified via polymerase chain reaction (PCR) using a technique similar to that described previously^1^. Reactions were conducted with 5 μL RNA and TaqMan Fast Virus 1-Step Master Mix (Applied Biosystems) with 500 nM primers and 200 nM probes. Primers and probes were as follows:

sgLeadSARSCoV2_F: 5’-CGATCTCTTGTAGATCTGTTCTC-3’

E_Sarbeco_P: 5’-FAM-ACACTAGCCATCCTTACTGCGCTTCG-BHQ1-3’

E_Sarbeco_R: 5’-ATATTGCAGCAGTACGCACACA-3’

Reactions were run on a QuantStudio 6 Pro Real-Time PCR System (Applied Biosystems) at the following conditions: 50 °C for 5 min, 95 °C for 20 sec, and 40 cycles of 95 °C for 15 sec and 60 °C for 1 min. Absolute quantification was performed in comparison to a standard curve. For the standard curve, the E subgenomic mRNA sequence was inserted into a pcDNA3.1 vector (Genscript) and transcribed using MEGAscript T7 Transcription Kit (Invitrogen) followed by MEGAclear Transcription Clean-Up Kit (Invitrogen). The lower limit of quantification was 50 copies.

### Histopathology

Histopathological analyses were performed as described previously^28^. Immunohistochemistry was used to visualize SARS-CoV-2 nucleocapsid antigen (rabbit polyclonal, GeneTex), and eosinophils by staining for eosinophil peroxidase (rabbit polyclonal, Atlas Antibodies). Inflammation was scored on the following scale based on % tissue affected: **0**, 0%; **1**, <10%; **2**, 10-25%; **3**, 26-50%; **4**, >50%. Viral antigen from IHC was scored on the following scale: **0**, minimal to absent; **1**, minimally abundant but clearly present; **2**, mildly abundant; **3**, moderately abundant; **4**, abundant. Eosinophils from IHC were scored on the following scale: **0**, within normal limits; **1**, minimal increase; **2**, mild increase; **3**, moderate increase; **4**, abundant.

### Statistics

All statistical analyses were performed in Prism (version 8.4, GraphPad). For comparisons between NHP vaccine groups at a single time point or dilution, a Kruskal-Wallis test with Dunn’s multiple comparisons test was performed. For comparisons over a time course, a 2-way ANOVA was employed with the Geisser-Greenhouse correction and the Tukey test to correct for multiple comparisons (Sidak’s test was used for comparing only two groups over a dilution series). For T cell assays a mixed-effect model with the Geisser-Greenhouse correction and a Tukey test was used, as not all samples were available at each time point. For comparisons of viral load in NHP and viral load and weight loss in hamsters over time, a 2-way ANOVA was employed with the Geisser-Greenhouse correction and a Dunnett test to correct for multiple comparisons, with comparisons to the PBS control group only. For comparisons of NHP titers post-challenge over time (not all samples were available at the last time point), a mixed-effect model was employed with the Geisser-Greenhouse correction and a Dunnett test, with comparisons controlled to the week 5 time point. For the weight loss versus binding titer correlation, a Spearman correlation test was performed. All binding titer and viral load data were log-transformed before performing statistical tests.

## Author Contributions

JRF, KSC, RAS, MGL, VL, TB, TMF, DC, RC, CG, MK, RvdM contributed to the concept or design of the study. JRF, BJF, KEF, ATN, APW, INM, MG, TSJ, CT, RLD, BF, SOC, SFA, EL, DRF, STN, MMD, ALZ, CA, SF, MJG, SS, VVE, KF, LL, HA, AC, AA, LP, and AVR collected and analyzed data. JPMT, AT, and EMC provided study coordination. ITT, TZ and DL contributed critical reagents. JRF, RAS, CB, SR were involved in the analysis and interpretation of the data. BFH, RAS, MSS, GA, MR, PDK, NJS, DCD, BSG provided supervision. All authors had full access to the data and approved the manuscript before it was submitted by the corresponding author

## Acknowledgement

The authors thank Jon Smith for coordinating the production and providing the vaccine antigens for the study, and Chris Case for project management.

