## Supplementary Figures for "Vaccination with SARS-CoV-2 Spike Protein and AS03 Adjuvant Induces Rapid Anamnestic Antibodies in the Lung and Protects Against Virus Challenge in Nonhuman Primates"

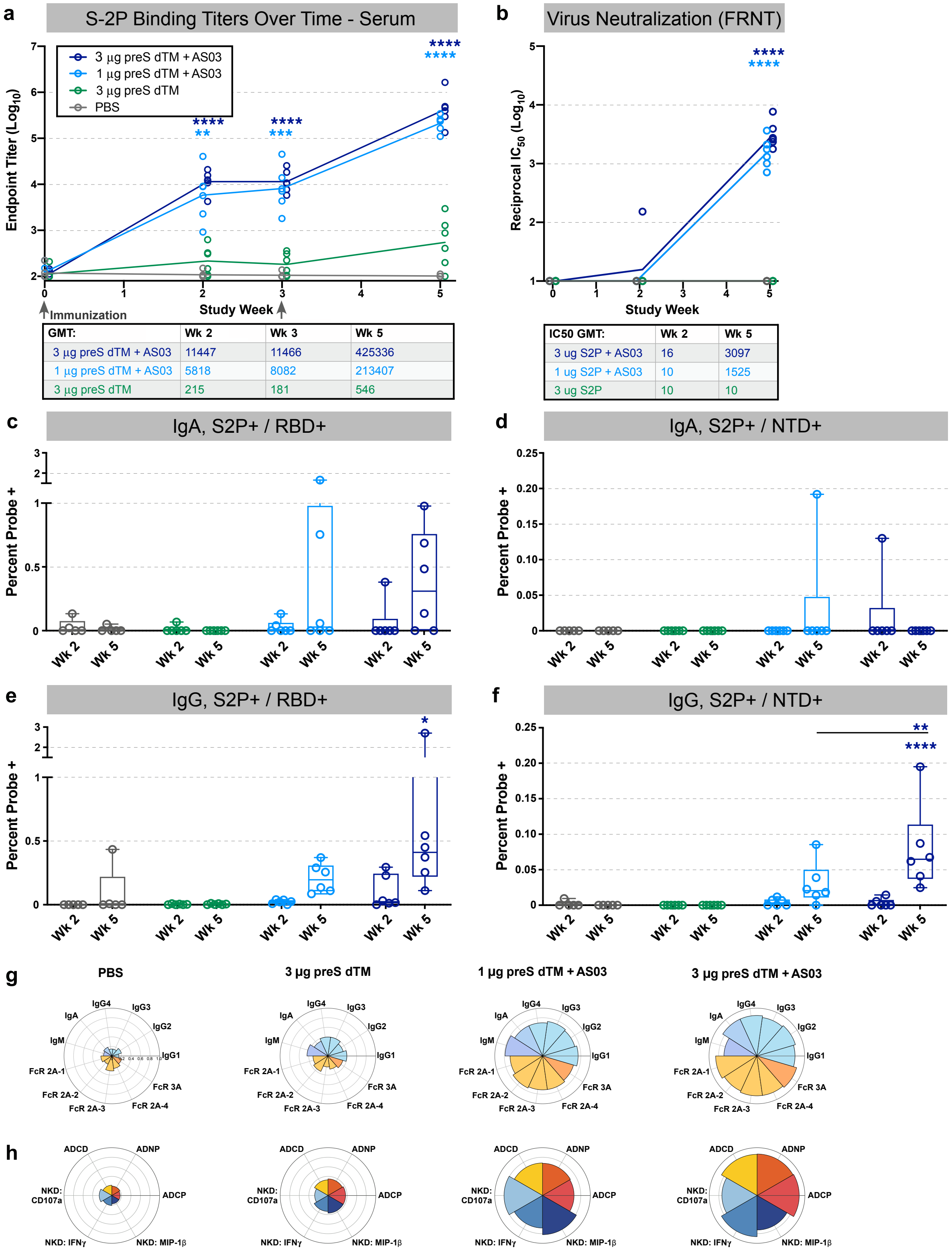


Supplementary Figure 1. AS03 adjuvant is required for functional antibody responses.
Rhesus macaques were immunized at weeks 0 and 3 with 1 or 3 µg of preS dTM adjuvanted with AS03 adjuvant or 3 µg of preS dTM formulated with PBS only. **a**, Endpoint binding titers over time following

vaccination. **b**, Live virus neutralization over time; $IC_{50}$ values are plotted. Symbols represent individual animals; bars and lines indicate group geometric means; geometric mean values for binding are indicated in table below. **c-f**, Antigen specific B cells were enumerated in the PBMC compartment using S-2P, RBD and NTD B cell probes after prime (week 2) and boost (week 5). IgA specific cells were found to bind both the S-2P and RBD probes **c**; and S-2P and NTD probes, **d**. IgG specific cells were found to bind both the S-2P and RBD probes, **e**; and S-2P and NTD probes, **f**. Symbols represent individual animals; box plots indicate the median and interquartile range; whiskers indicate minimum and maximum data points. **g-h,** A systems serology approach was used to measure antibody subtypes, isotypes and $F_C$ receptor ($F_CR$) binding (**g**), as well as $F_C$-mediated effector functions (**f**) using spike protein. Flower plots indicate a normalized Z-score for each parameter. ADCD, antibody dependent complement deposition; ADNP, antibody dependent neutrophil phagocytosis; ADCP, antibody dependent cellular phagocytosis; NKD, natural killer cell degranulation. Asterisks indicate significance compared to the PBS control group at each time point (unless otherwise indicated): *, $p<0.05$; **, $p<0.01$; ***, $p<0.001$ ****, $p<0.0001$.

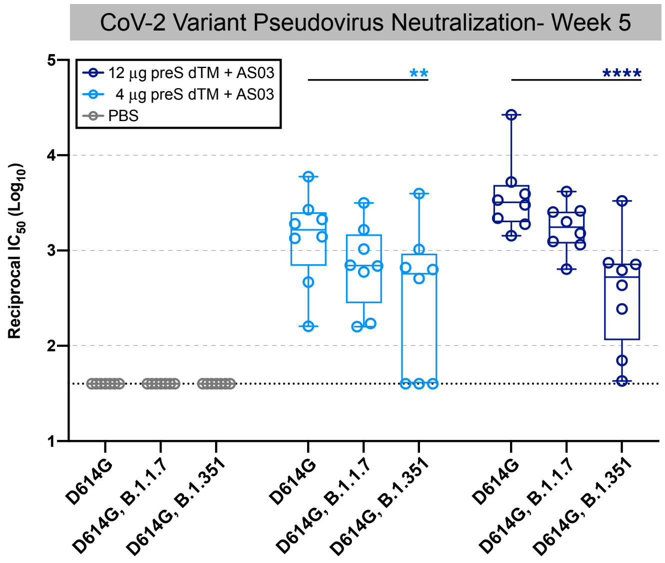


| Endpoint GMT: | D614G | D614G, B.1.1.7 | D614G, B.1.351 |
|---|---|---|---|
| 12 µg preSΔTM + AS03 | 3671 | 1719 | 370 |
| 4 µg preSΔTM + AS03 | 1319 | 674 | 294 |

Supplementary Figure 2. Pseudoviral neutralization of SARS-CoV-2 variants.
Pseudoviral neutralization responses in rhesus macaques immunized with 4 or 12 ug of preS dTM adjuvanted with AS03 adjuvant at weeks 0 and 3. Plasma from week 5 were tested against the parental D614G virus, or against B.1.1.7 (UK variant) or B.1.351 (South African variant) also containing the D614G mutation. Geometric mean values for neutralization are indicated in the table below the graph. Symbols represent individual animals; box plots indicate the median and interquartile range; whiskers indicate minimum and maximum data points. Asterisks indicate significance compared to the D614G virus for each vaccine group. **, $p<0.01$; ****, $p<0.0001$.

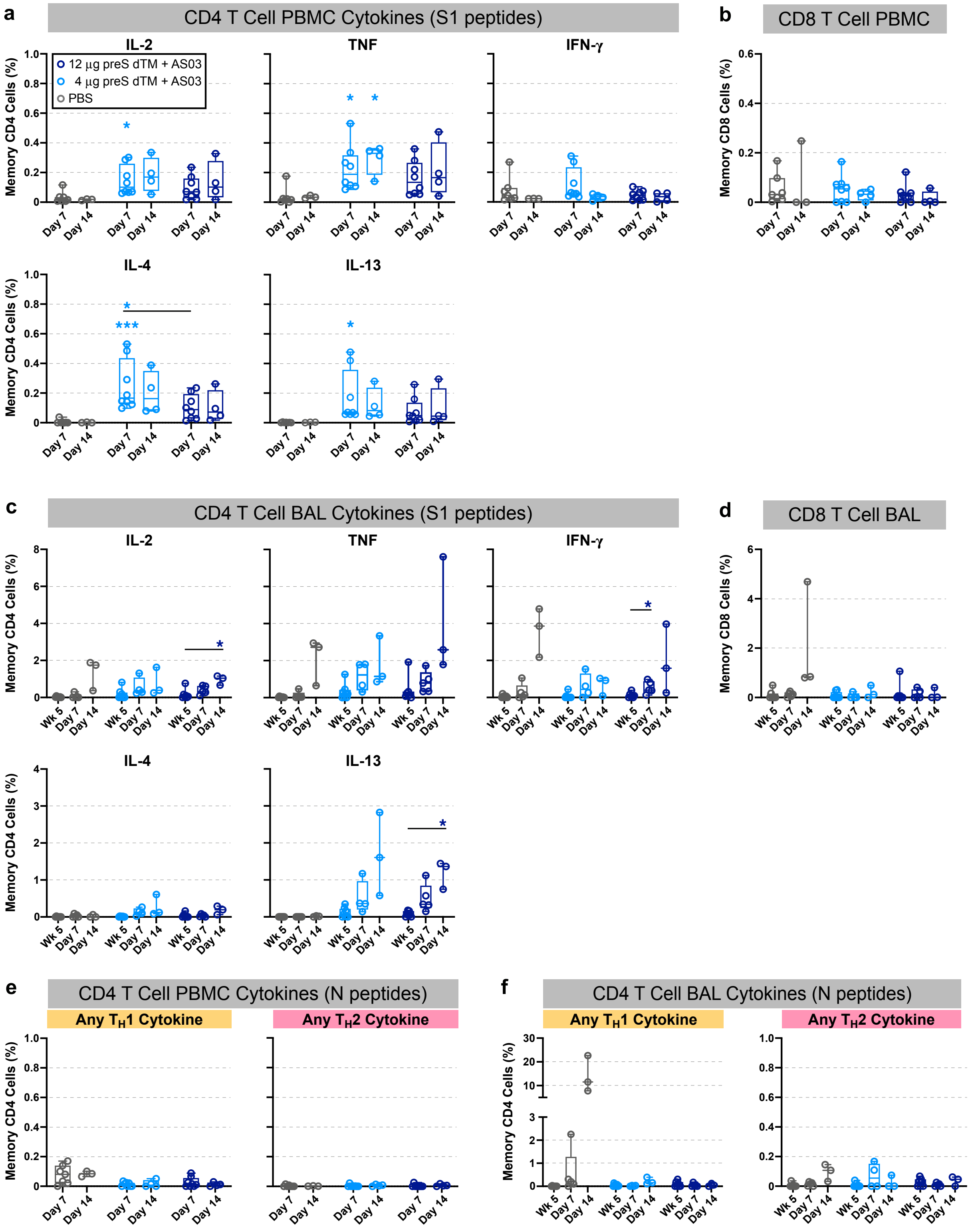


Supplementary Figure 3. T cell responses following SARS-CoV-2 challenge.
PBMC and bronchoalveolar lavage (BAL) cells were collected prior to challenge (week 5) and on days 7 and 14 following SARS-CoV-2 challenge. Cells were stimulated with pools of peptides covering the

spike (S1) or nucleocapsid (N) proteins. **a**, Percent of PBMC memory CD4 T cells making the indicated cytokines following stimulation with S1 peptides. **b**, Percent of memory PBMC CD8 T cells making any $T_H1$ cytokine following stimulation with S1 peptides. **c**, Percent of memory BAL CD4 T cells making the indicated cytokines following stimulation with S1 peptides. **d**, Percent of BAL memory CD8 T cells making any $T_H1$ cytokine following stimulation with S1 peptides. **e**, Percent of PBMC memory CD4 T cells making any $T_H1$ (left graph) or any $T_H2$ cytokine (right graph) following stimulation with N peptides. **f**, Percent of BAL memory CD4 T cells making any $T_H1$ (left graph) or any $T_H2$ cytokine (right graph) following stimulation with N peptides. Symbols represent individual animals; box plots indicate the median and interquartile range; whiskers indicate minimum and maximum data points. Asterisks indicate significance compared to the PBS control group for each time point (unless otherwise indicated) as follows: *, $p<0.05$; ***, $p<0.001$.

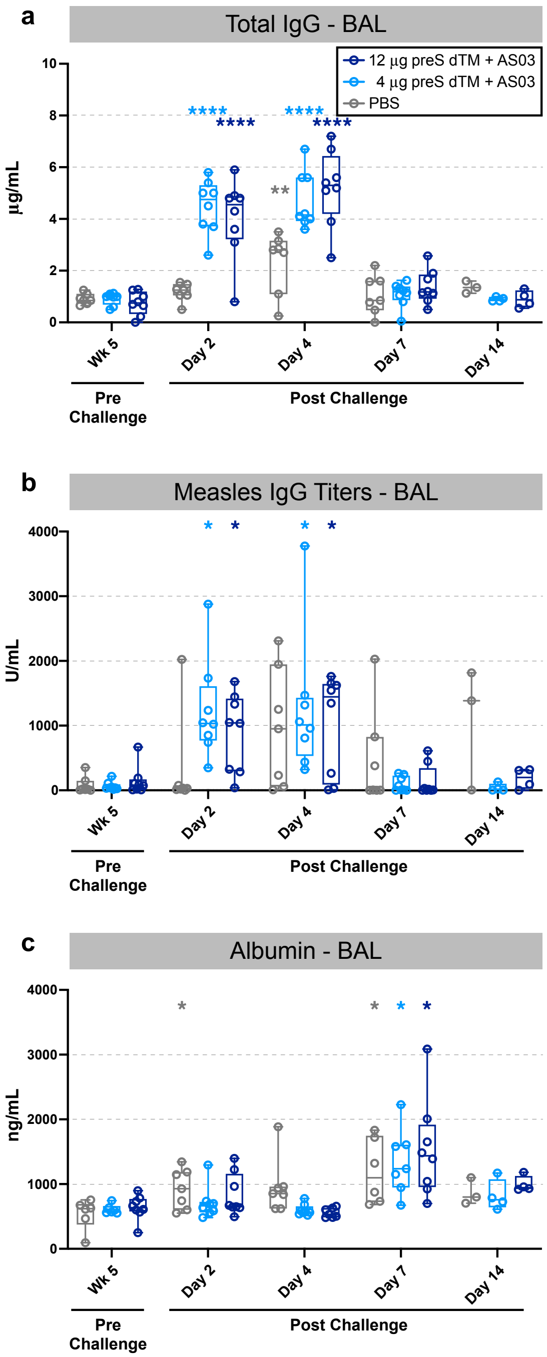


Supplementary Figure 4. Nonspecific antibody responses in the lung following SARS-CoV-2 challenge. Bronchoalveolar lavage (BAL) supernatant was collected prior to challenge (week 5) and on days 2, 4, 7 and 14 following SARS-CoV-2 challenge. **a**, Total IgG concentration titers in BAL. **b**, Measles responses in BAL. **c**, Albumin levels in BAL. Symbols represent individual animals; box plots indicate the median and interquartile range; whiskers indicate minimum and maximum data points. Asterisks indicate significance compared to the week 5 time point as follows: *, p<0.05; **, p<0.01; ****, p<0.0001.

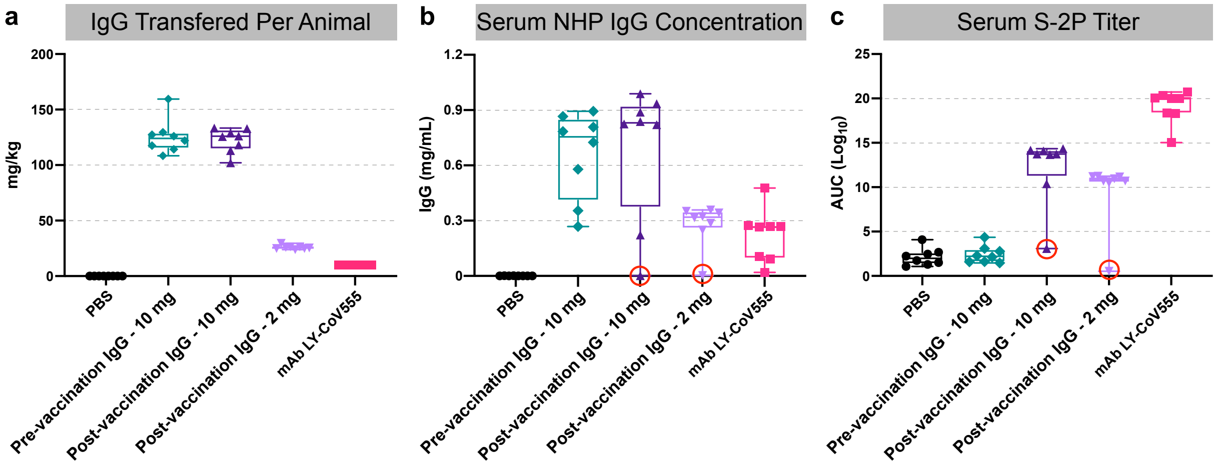


Supplementary Figure 5. Passive transfer of IgG from vaccinated NHP to hamsters.
Total IgG was isolated from rhesus macaques immunized with 3 µg of AS03-adjuvanted preS dTM. 10 or 2 mg total IgG was transferred to hamsters, or PBS was given as a negative control; serum was collected one day later, on the day of challenge. **a**, The amount of IgG transferred based on the weight of each animal; mAb 555 was delivered at 10 mg/kg. **b**, NHP total IgG ELISA was used to measure the concentration of NHP IgG in hamster serum, by extrapolating from an NHP IgG standard curve. **c**, S-2P ELISA reactivity of hamster serum shown as area under the curve. Red circles indicate 2 animals that were improperly infused and were thus excluded from the weight loss analysis in Figure 6b. Symbols represent individual animals; box plots indicate the median and interquartile range; whiskers indicate minimum and maximum data points.

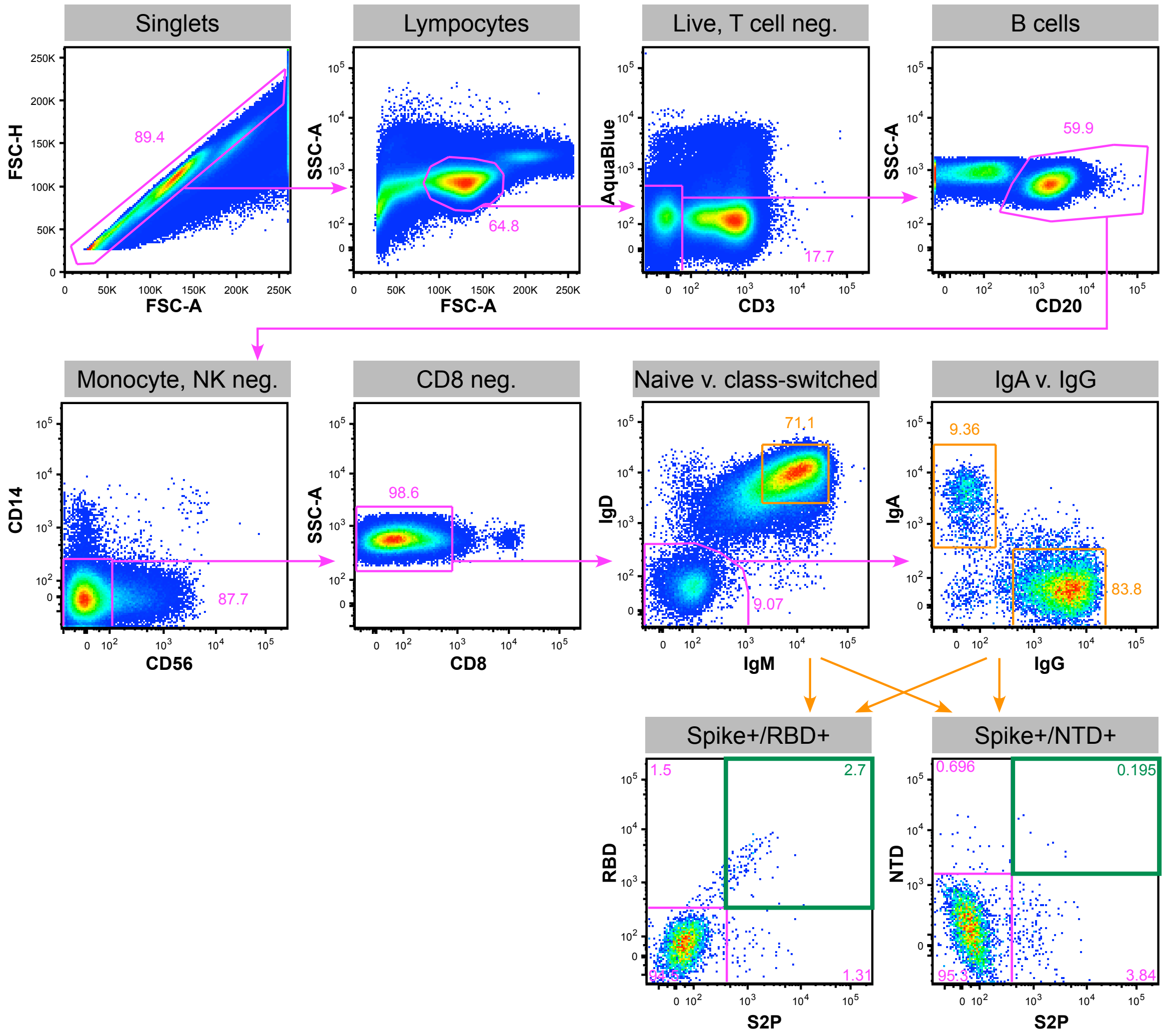


Supplementary Figure 6. Gating tree for isotyping antigen-specific B cells.
Vaccine-specific B cells were identified from PBMC by first gating on singlets, followed by gating for lymphocytes based on forward and side scatter. B cells were then negatively gated for CD3, positively gated on CD20, then negatively gated on CD56, CD14 and CD8. Class-switched (memory) B cells were gated as IgD and IgM negative; double-positive cells in this plot were gated as naïve B cells. Memory B cells were then further separated by IgA and IgG markers, followed by gating for SARS-CoV-2 spike antigen-specific cells by gating for cells that were positive for both S-2P and RBD, or S-2P and NTD probes.
